# Spatially Varying Trophic Effects of Reservoir-Derived Plankton on Stream Macroinvertebrates Among Heterogeneous Habitats Within Reaches

**DOI:** 10.1101/870717

**Authors:** Shinji Takahashi, Yasuhiro Takemon, Tatsuo Omura, Kozo Watanabe

## Abstract

1. Dam reservoirs often supply high amounts of plankton to downstream reaches, leading to a critical shift of trophic origins of stream ecosystems from natural sources (e.g. attached algae and terrestrial inputs) to reservoir-oriented plankton. Although this is a widely observed phenomenon, previous studies focused only on lotic habitats (e.g. riffles) rather than lentic habitats such as backwaters and isolated ponds (IP).
2. Using a stable isotope three-source mixing model, we evaluated trophic contributions of reservoir-derived plankton, epilithon and terrestrial leaves to stream macroinvertebrates at four dam outlet reaches and two reference reaches in the Natori River catchment, Japan. We compared four different habitat types co-occurring within the reaches: lotic habitat (riffle and pool), bar-head (BH) lentic habitat, bar-tail (BT) lentic habitat (backwater) and isolated pond (IP) on sandy bars.
3. The trophic contributions of reservoir-derived plankton were significantly lower in lentic habitats (BH, 15.4%; BT, 10.4%; IP, 9.1%) than in lotic habitats (mean, 27.7%). This was especially notable for filter feeders that feed on suspended fine particulate organic matter (SFPOM). The three-source model analysis indicated a lower biomass proportion of dam plankton in lentic SFPOM (mean, 21.2%) than in lotic SFPOM (mean, 35.6%). This difference in SFPOM composition was reflected in the lower trophic contribution of dam plankton to lentic filter feeders.
4. The abundance ratio of filter feeders in the community was decreased in lentic habitats, while the abundance ratios of collector-gatherers, scrapers and shredders were increased. Macroinvertebrates in lentic habitats fed on sources less mixed with reservoir-derived plankton (e.g. benthic coarse particulate organic matter [BCPOM], benthic fine particulate organic matter [BFPOM] and epilithon); therefore, the trophic impact of reservoirs was indistinctive at the community level, indicating that lentic habitats can function as trophic refugia to mitigate the trophic impact of reservoirs.
5. Because lentic habitats were decreased in area (accounting for 5.7% of average total area) in the downstream reaches of dams due to riverbed degradation, lentic habitats must be created in order to restore the trophic impact of reservoirs in river ecosystems.

## Introduction

A key question in the field of food web ecology is how spatial environmental heterogeneity drives diverse food resources and food web structures within a limited space (Wissel & Fry, 2005; Leigh & Sheldon, 2009; Sereda et al., 2012; Kaymak et al., 2018). Riverine reach is a typical template that has large spatial variations in geomorphologic and hydrologic conditions among habitat patches (e.g., riffle, pool, backwater and pond) (Yarnell et al., 2006; Heino, 2013). These morpho-hydrologic variations may influence the patterns of material and energy flow among habitats within reaches, sustaining spatial heterogeneity in the abundance and composition of food sources (Wanner et al., 2002). Assemblage trophic structure varies spatially in association with longitudinal and lateral gradients of geomorphology, environmental conditions and disturbance regimes in rivers (Hoeinghaus et al., 2007; East et al., 2017). Some habitats include highly stored terrestrial detritus and epilithon in backwaters and IPs on sandy bars (Nakajima et al., 2006; Flinn et al., 2008), abundant coarse particulate organic matter deposited in riparian ponds (Langhans et al., 2013), high biomass of phytoplankton in water bodies of medium age (2 to 10 days) (Hein et al., 2003) and high biomass of periphyton in lentic habitats (Biggs & Close, 1989). However, little is known about their ecological consequences for food web structure and energy flow at the community’s consumer levels.

Dam reservoirs may artificially alter the spatial distribution of food sources and community food web structure at dam outlet reaches (e.g. Hoffsten, 1999; Doi et al., 2008; Helmus et al., 2013; Wellard Kelly et al., 2013; Martinez et al., 2013; Murphy et al., 2017; Four et al., 2019). A typical phenomenon often observed below dams is an increased supply of fine particulate organic matter (FPOM) as a consequence of high primary production in reservoirs, which is especially notable in eutrophic lakes (Voelz & Ward, 1996). The high load of FPOM sometimes leads to a critical shift of the main trophic origins of stream ecosystems from natural sources (e.g. attached algae and terrestrial particulate organic matter) to reservoir-derived plankton and also a shift of the functional feeding group (FFG) composition of macroinvertebrates to a structure more dominated by filter feeders (Sheldon and Oswood, 1997; Kobayashi et al., 2011).

Although the trophic influence of dam plankton on downstream consumers has been widely tested and validated, previous tests were conducted only in lotic habitats or riffles (e.g. Watanabe & Omura, 2007; Doi et al., 2008; Tagliaferro et al., 2013). To our knowledge, the trophic effect of dams on lentic habitats, such as backwaters and IPs on sandy bars or on riparian terraces, has not been tested (Malard et al., 2002; Takemon, 2007). Therefore, supplementary tests that compare different habitat types in dam outlet reaches may provide better insights into the role of spatial environmental heterogeneity in the formation of spatially varying trophic structures.

Carbon and nitrogen stable isotope analysis is a widely used approach to estimate trophic contributions of different potential food sources to aquatic animals (Phillips et al., 2005; Leberfinger et al., 2011), assuming a monotonic change of isotopic properties through the anabolic process (Rounick et al., 1982; Reid et al., 2008). Autochthonous (e.g. epilithon) and allochthonous (e.g. terrestrial litter) organic matter are the two main sources of trophic contributions in river ecosystems. They have different isotopic signatures due to their different photosynthetic mechanisms and activities (Finlay et al., 2002). In dam outlet reaches, lentic plankton produced in reservoirs could be an additional food source. Dam plankton tends to show distinct isotopic signatures (i.e. low carbon and high nitrogen isotopic signatures) compared with the *in situ* sources produced in rivers (Ock & Takemon, 2014). Based on the reservoir-specific isotopic properties of plankton, several isotopic studies found high trophic contributions of drifting dam plankton to downstream macroinvertebrate communities (Richardson & Mackay, 1991; Monaghan et al., 2001; Watanabe & Omura, 2007; Doi et al., 2008; Mercado-Silva et al., 2009). However, these previous tests were conducted only in riffles, and spatial heterogeneity among different habitat types was out of their focus (but see Ock & Takemon (2014) for the difference between riffle and pool).

In this study, using a carbon and nitrogen stable isotope mixing model (Finlay et al., 2002; Reid et al., 2008; Molina et al., 2011; Kominoski et al., 2012), we evaluated the trophic contributions of reservoir-derived plankton, epilithon and leaves to stream macroinvertebrates at four dam outlet reaches in the Natori River catchment, northeastern Japan, along with two reference reaches. We compared the trophic contributions of reservoir-derived plankton among four different habitat types co-occurring within the reaches. Our hypothesis was that differences among habitats in the composition and abundance of food sources induce spatially heterogeneous patterns in the trophic contributions of reservoir-derived plankton. The trophic effect of dam plankton on macroinvertebrates in lentic habitats may be mitigated by low rates of migration of dam plankton and/or accumulation of organic matter of stream origin. By combining data matrices obtained through stable isotope, ecological and GPS analyses, we found overall significant results that support our hypothesis.

## Methods

### Study sites

Field surveys and sample collection were conducted in six reaches in two basins in the Natori River catchment in Miyagi Prefecture, northeastern Japan (Fig. S1, Supporting Information). Each basin has one dam in the upstream section, the Kamafusa (K) and Ohkura (O) dams. Four reaches were selected from the downstream (D) reaches of dams (KD1, KD2, KD3 and OD), and two reaches were selected from upstream (U) reaches of the same dams as references (KU and OU) (i.e. no discharge from the dams). The three reaches below the Kamafusa dam are longitudinally located along a corridor with different water distances of 0.6 km (KD1), 2.7 km (KD2) and 6.3 km (KD3) from the dam. The Ohkura dam has one downstream study reach (OD) located 2.0 km below the dam. Although there is a small mountain runoff flow into the downstream reaches of the dams, most of the water in the study reaches is discharged from the dam reservoirs. The primary land use types around the six study reaches are agricultural areas and wasteland, and the channels have a complete open canopy. Only OU located in the forest area and the channel is covered by canopy. The Kamafusa and Ohkura dams have 45.5 and 82.0 m vertical and 177.0 and 323.0 m wide lengths and water storage capacities of 3.9 × 10^7^ and 2.5 × 10^7^ m^3^, respectively. High concentrations of phytoplankton are often observed in both dam reservoirs, especially during early summer (June and July).

We investigated spatial habitat structure in the six reaches once for each reach at KU and KD2 on 2 July 2008 and at KD1, KD3, OU and OD on 2 October 2008. We identified up to four habitat types for each reach: lotic (riffle and pool), bar-head (BH) wando, bar-tail (BT) wando (backwater) and isolated pond (IP) on sandy bars using a high-precision GPS (ProMark3; Thales, France). Geographic positions were recorded at 2-s intervals while walking along contours of the habitat and later post-corrected using base station data to obtain a precision of 0.1 m or less. Water surface area, frequency and shape complexity of each habitat type in a reach were calculated using ImageJ v.1.42 software (NIH, Bethesda, MD, USA). In the field, we also measured current water velocity and water depth in each habitat type using a current metre (VP-201; Kenek, Tokyo, Japan) and a ruler with five replicates per habitat type per reach.

We collected quantitative samples of macroinvertebrate communities from each habitat type found in the reaches using a Surber sampler (30 × 30 cm, 250-μm mesh) with three replicates per habitat type per reach once during July and October 2008. The samples were preserved in 99.5% ethanol and separated from detritus and sediment debris in the laboratory under a microscope with 150× magnification. Organisms were identified to the lowest taxonomic level possible (mostly at the species level) based on Kawai and Tanida (2005) and were assigned to one of five FFGs (filter feeders, collector-gatherers, scrapers, shredders and predators) based on the classification of Takemon (2005) modified from Merritt and Cummins (1996) for Japanese freshwater macroinvertebrates.

Epilithon (dominated by attached algae), suspended fine particulate organic matter (SFPOM) (< 1.0 mm), benthic fine particulate organic matter (BFPOM), benthic coarse particulate organic matter (> 1.0 mm) (BCPOM) and leaves were collected from three randomly selected locations per habitat type per reach on the same day of collection of macroinvertebrates. Epilithon was collected from stones using a toothbrush, washed in pure water, and filtered onto precombusted Whatman GF/F glass filters (0.7-µm nominal pore size). SFPOM was collected by filtering 1 to 4 litres of surface river water through a 1.0-mm sieve onto a GF/F filter. BFPOM was collected by placing a plastic tube sampler (diameter, 25 cm; depth, 50 cm) on the riverbed, disturbing the benthic material in the sampler by hand, collecting the turbid water and processing the sample by the same pretreatment as for SFPOM. Leaves were collected from several different plant species in each reach. BCPOM was collected using Surber nets together with the macroinvertebrates. Plankton was collected from the reservoirs at three levels (0, 3 and 10 m from the water surface) near the water intakes of the dams and processed by the same pretreatment as for SFPOM in river water. Finally, nine replicates of the epilithon, SFPOM, BFPOM, BCPOM and dam plankton samples were prepared for carbon (n = 3) and nitrogen (n = 3) isotope analyses and for ash-free dry mass (AFDM) analysis (n = 3) (Flinn, 2008), and six replicates of leaf samples were prepared for carbon (n = 3) and nitrogen (n = 3) isotope analysis. All samples were stored at −20°C prior to further processing.

### Stable isotope analysis

For stable isotope analysis, macroinvertebrates and potential food sources were acidified with 1 mol l^−1^ HCl to remove carbonate, and the remaining material was rinsed with distilled water and kept in a freezer (Walters et al., 2007). All samples were freeze-dried and homogenised before stable isotope analysis. The samples, ranging from 0.5 to 1.5 mg dry weight for macroinvertebrates and from 1 to 20 mg dry weight for the potential food sources, were weighted into tin capsules. Carbon and nitrogen isotope ratios (^13^C/^12^C and ^15^N/^14^N) were measured using an elemental analyser (NA2500; CE Instruments, USA) coupled to a continuous flow mass spectrometer (Finnigan MAT, Delta Plus; Thermo Fisher, USA). Stable isotope ratios were evaluated in δ notation as the deviation from standards (Pee Dee belemnite for δ^13^C and atmospheric nitrogen for δ^15^N), calculated as δ^13^C or δ^15^N = [(R_sample_/R_standard_)−1] ×10^3^, where R is ^13^C/^12^C or ^15^N/^14^N, respectively. Typical precision of the analyses was ± 0.5‰ for δ^15^N and ± 0.2‰ for δ^13^C.

A Bayesian mixing model of stable isotope analysis in R (SIAR) (Parnell et al., 2010) was used to calculate the relative contribution of each potential food source to the diet of macroinvertebrates. The potential food sources used in the three-source model in the dam outlet reaches were epilithon, leaves and dam plankton. The carbon and nitrogen isotope enrichments in the model were set to + 0.4‰ (McCutchan et al., 2003) and 3.4‰ (Post, 2002), respectively. Before performing the model calculations, we confirmed significant differences in mean isotopic values among the three potential food sources using one-way analysis of variance (ANOVA). In the two upstream reaches of dams (KU and OU), we alternatively used a two-source mixing model (Peterson & Fry, 1987; Molina et al., 2011) to determine the relative contribution of epilithon to the diet of consumers (*f*) against leaves: *f* =[(δ^13^Cconsumer–δ^13^Cleaves) / (δ^13^Cepilithon–δ^13^Cleaves)] x 100, where δ^13^Cconsumer is the δ^13^C of each consumer taxon, and δ^13^C_epilithon_ and δ^13^C_leaves_ are the δ^13^C values of epilithon and terrestrial plants. When *f* was less than 0 in the calculation, 0 was used as *f*.

### Statistical analysis

The mean levels of hydraulic variables (depth and velocity), amounts of the potential food sources (epilithon, SFPOM, BFPOM and BCPOM), trophic contributions of the three sources (dam plankton, epilithon and leaves) to whole macroinvertebrate communities and FFGs, and biomass proportions of the three sources in SFPOM, BFPOM and BCPOM were compared among the four habitat types (lotic, BH, BT and IP) using one-way ANOVA followed by the Tukey–Kramer test for multiple pairwise comparisons. The trophic contributions of dam plankton were also compared among FFGs for each habitat type. Simple correlations between the trophic contributions of dam plankton to macroinvertebrate communities and FFGs and the biomass proportions of dam plankton in SFPOM, BFPOM and BCPOM were tested by ANOVA. All statistical analyses were carried out using SPSS Statistics v.17.0 (SPSS, Chicago, IL, USA).

## Results

### Habitat structure, hydraulic and food source conditions

Lotic and BH habitats commonly occurred throughout the six study reaches. BT occurred at KU, KD3 and OD, and IP occurred at KD1 and KD3. IP occurred only at dam outlet reaches. Throughout the six reaches, lotic habitat always had the highest proportion of water area in the reaches (88.5% to 96.8%), while lentic habitats (BH, BT and IP) accounted for a minor proportion of water area (1.4% to 11.5%) (Table 1).

**Table 1.**
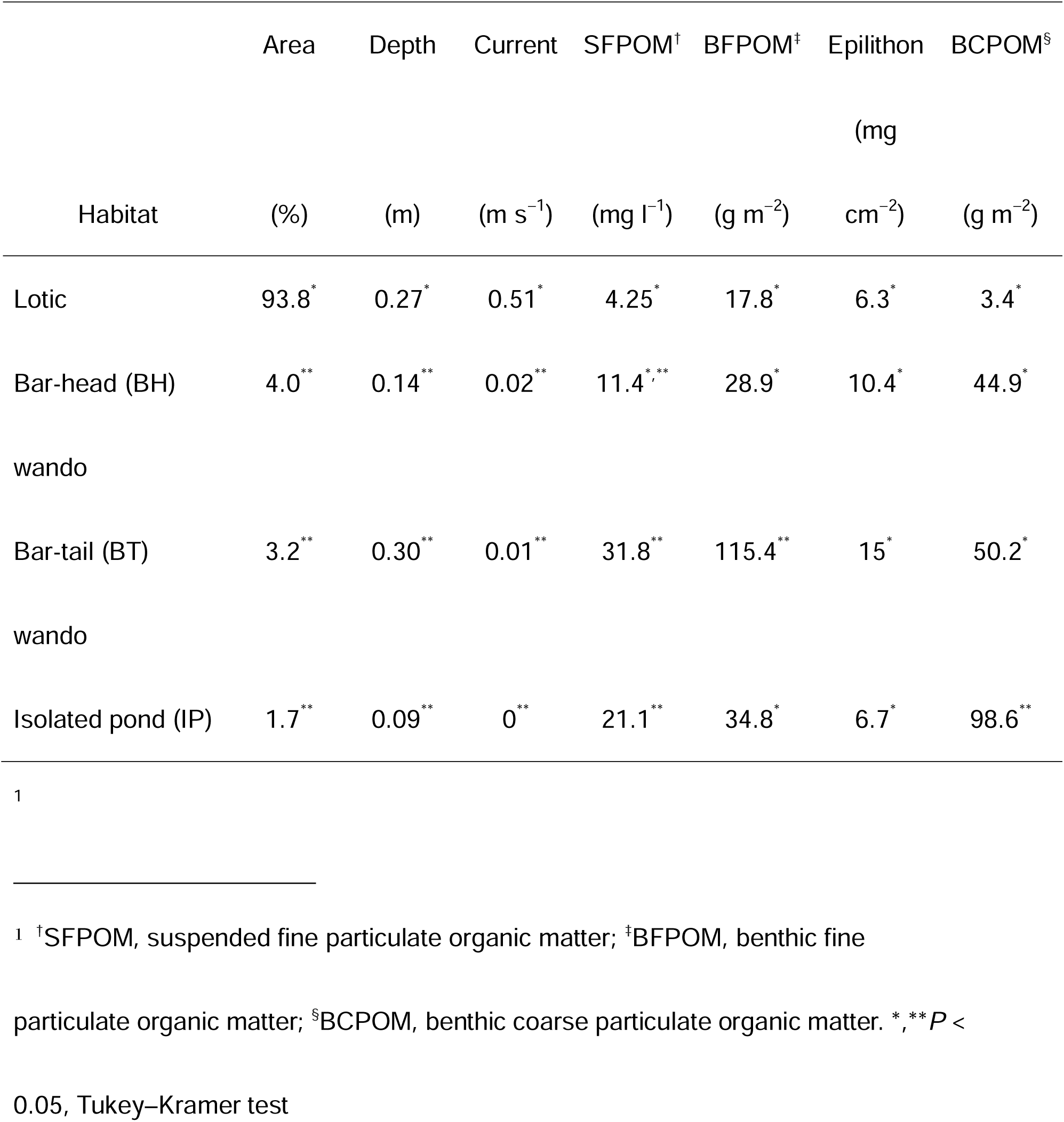
Mean values of hydraulic variables and amounts of potential food sources in each habitat type among six study reaches

Mean water depth (F_3, 166_ = 24.7, *P* < 0.001) and mean current velocity (*F*_*3, 164*_ = 105.5, *P* < 0.001) (ANOVA) differed significantly among the four habitat types (Table 1). Multiple pairwise comparison tests found significantly higher water depths in the BT (mean, 0.30 m) and the lotic habitat (mean, 0.27 m) than in the BH (0.14 m) and IP (0.09 m) (*P* < 0.001, Tukey–Kramer test). The mean current velocity was significantly higher in the lotic habitat (0.51 m s^−1^) than in the three lentic habitat types (BH, 0.02 m s^−1^; BT, 0.01 m s^−1^; IP, 0.0 m s^−1^; *P* < 0.001, Tukey–Kramer test).

Differences among habitats in the availability of food sources were also evident. The mean biomass (AFDM) of potential food sources was significantly different among the four habitat types (SFPOM, *F*_3, 91_ = 7.554, *P* < 0.001; BFPOM, *F*_3,82_ = 19.421, *P* < 0.001; BCPOM, *F*_3, 58_ = 11.717, *P* < 0.001), except for epilithon (*F*_3, 73_ = 2.241, *P* > 0.05). Multiple pairwise comparison tests found significantly larger mean biomasses of SFPOM, BFPOM and BCPOM in the three lentic habitat types than in the lotic habitat (*P* < 0.01, Tukey–Kramer test). SFPOM biomass was higher in IP and BT than in the lotic habitat (*P* < 0.01, Tukey–Kramer test), and BH was those in between (*P* > 0.05, Tukey–Kramer test). BFPOM biomass was highest in the BT (*P* < 0.001, Tukey–Kramer test), and BCPOM biomass was highest in the IP (*P* < 0.01, Tukey–Kramer test). Three habitat types, but not the IP, showed significant increases in mean SFPOM biomass from the upper to the lower reaches of the dam (lotic, from 1.3 to 5.8 mg l^−1^; BH, from 2.3 to 16.8 mg l^−1^; BT, from 2.3 to 35.5 mg l^−1^; *P* < 0.05, *t-*test). BFPOM biomass increased only in the BT (from 37.9 to 154.2 mg l^−1^, *P* < 0.01), and epilithon and BCPOM biomass did not change between the upper and lower reaches of the dam.

### Macroinvertebrate community

We collected a total of 5547 macroinvertebrates and identified 125 taxa from the 17 habitats among the six reaches (Table 2). The majority of taxa were of the orders Trichoptera (40 taxa), Ephemeroptera (40 taxa), Plecoptera (11 taxa) and Diptera (11 taxa). Taxon richness ranged widely from 11 to 42 taxa among the 17 habitats, with significant differences between the lotic (mean, 32.7) and lentic habitat types (IP, 16.0; BT, 16.7; BH, 20.0; *P* < 0.05, Tukey–Kramer test). Total abundance (N) and total biomass (W) were also higher in the lotic habitat than in the lentic habitat types (*P* < 0.01, *t-*test), except for BH and BT at KU, where C*hironomus* sp. (70 and 921 individuals/0.27 m^2^, respectively) were highly abundant, and IP at KB3, where *Cloeon dipterum* (967 individuals/0.27 m^2^) was highly abundant. Among the 125 taxa, 37 and 39 taxa occurred only in lentic or lotic habitats, respectively. *Antocha* sp. were commonly found throughout all 17 habitats with high abundance. The most frequently occurring taxa in the lotic habitats were *Macrostemum radiatum, Cincticostella okumai* and *Baetiella* sp., while *Hydropsyche setensis* occurred throughout all six lotic habitats.

**Table 2.**
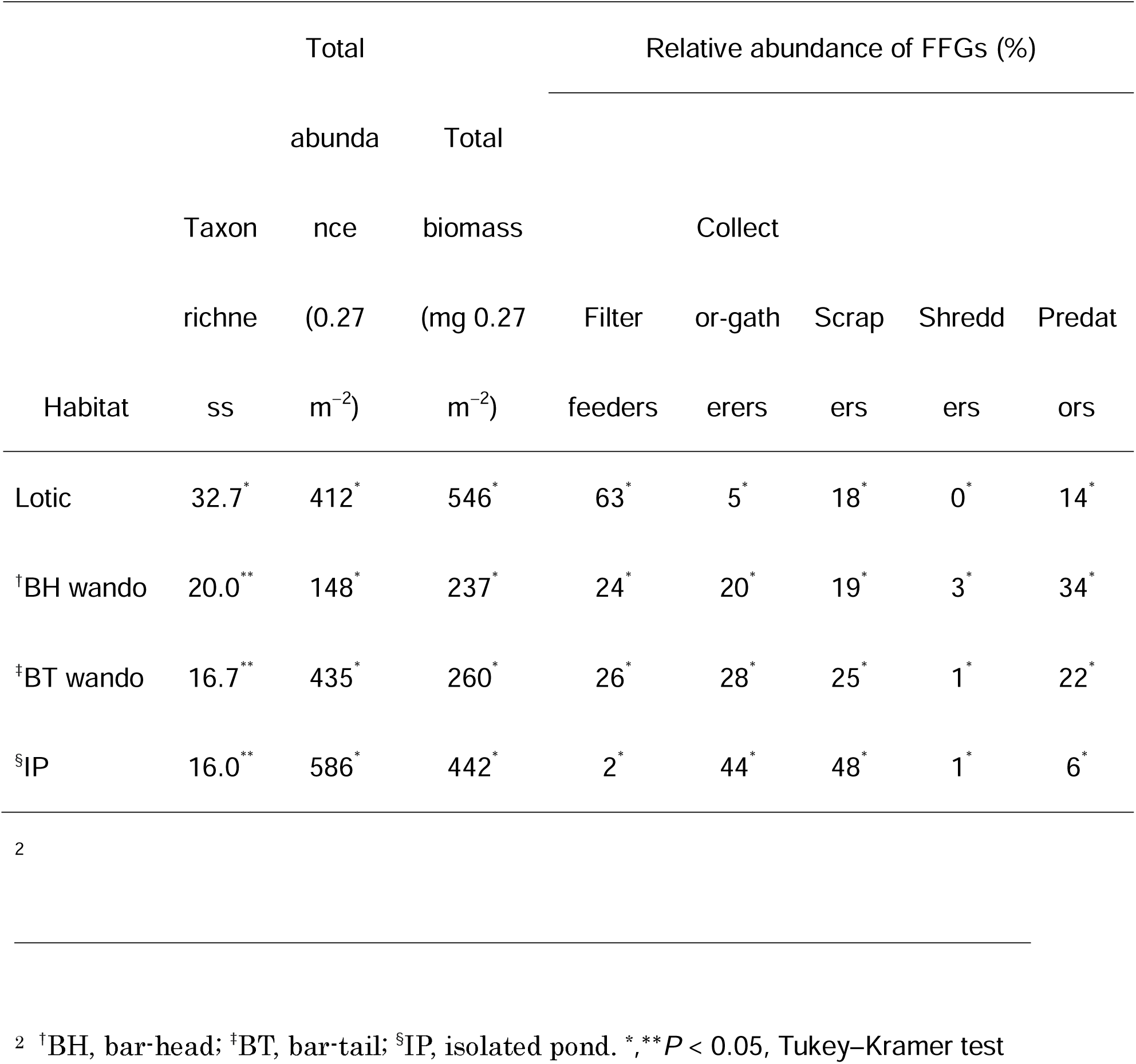
Mean values of species richness, total abundance and total biomass of macroinvertebrate communities and percentages of functional feeding groups (FFGs) abundance in each habitat type among six study reaches

In the dam downstream reaches, the mean proportional abundance of filter feeders in the macroinvertebrate community was significantly higher in lotic habitats than in lentic habitats (*F*_1, 13_ = 24.3; *P* < 0.001, two-way ANOVA), whereas this was not true for the other FFGs. The spatial variation in the proportion of filter feeders among the six lotic habitats was significantly and positively correlated with the variation in SFPOM concentration (*r* = 0.954, *P* = 0.003). We did not detect any other significant correlations between the proportion of FFGs and the biomass of SFPOM, BFPOM and BCPOM in any habitat type.

### Carbon and nitrogen Isotope signatures

The carbon (δ^13^C) and nitrogen (δ^15^N) isotope signatures varied among potential food sources in all habitat types within reaches (*P* < 0.01, ANOVA). Epilithon had the highest δ^13^C (–22.4 ± 3.4‰) and δ^15^N (4.1 ± 2.3‰) values, while leaves had the lowest δ^13^C (–29.8 ± 1.4 ‰) and δ^15^N (0.5 ± 0.2‰) values in each habitat. Dam plankton collected from the reservoirs had low δ^13^C (–28.2 ± 0.5‰) and high δ^15^N (4.9 ± 0.7‰) values. In the dam outlet reaches, δ^13^C and δ^15^N values for SFPOM and BFPOM fell within the range of values for epilithon, leaves and dam plankton, allowing the three-source mixing model to be run. All three lentic habitat types had significantly lower mean proportions of dam plankton in SFPOM than the lotic habitat (*P* < 0.05, Tukey–Kramer test) (Table 3). On the other hand, the mean proportion of epilithon in SFPOM was higher in the BH and IP than in the lotic habitat (*P* < 0.01,Tukey–Kramer test). BFPOM had higher proportions of dam plankton only in the IP (*P* < 0.05, Tukey–Kramer test). In upstream reaches of dams, there were no significant differences among habitat types in the proportion of the three sources (epilithon, leaves and dam plankton) in SFPOM and BFPOM.

**Table 3.** Comparison of mean percentages of trophic contributions of potential food sources in SFPOM, BFPOM and the macroinvertebrate community among four habitat types, calculated by two- or three-source mixing models

| Habitat | Reference |  | Dam downstream |  |  |
| --- | --- | --- | --- | --- | --- |
|  | Epilithon | Leaves | Plankton | Epilithon | Leaves |
| <sup>†</sup>SFPOM |  |  |  |  |  |
| Lotic | 67.0<sup>*</sup> | 33.0<sup>*</sup> | 35.6<sup>*</sup> | 34.1<sup>*</sup> | 30.3<sup>*,**</sup> |
| <sup>§</sup>BH wando | 55.7<sup>*</sup> | 44.3<sup>*</sup> | 23.8<sup>*</sup> | 51.5<sup>*</sup> | 24.9<sup>*</sup> |
| <sup>¶</sup>BT wando | 50.4<sup>*</sup> | 49.6<sup>*</sup> | 20.5<sup>**</sup> | 37.8<sup>*,**</sup> | 41.7<sup>**</sup> |
| <sup>∞</sup>IP | — | — | 19.3<sup>**</sup> | 54.0<sup>**</sup> | 26.8<sup>*,**</sup> |
| <sup>‡</sup>BFPOM |  |  |  |  |  |
| Lotic | 65.3<sup>*</sup> | 34.7<sup>*</sup> | 22.1<sup>*</sup> | 41.6<sup>*</sup> | 36.3<sup>*</sup> |
| <sup>§</sup>BH wando | 53.7<sup>*</sup> | 46.3<sup>*</sup> | 17.9<sup>*</sup> | 46.3<sup>*</sup> | 35.7<sup>*</sup> |
| <sup>¶</sup>BT wando | 56.4<sup>*</sup> | 43.6<sup>*</sup> | 17.7<sup>*</sup> | 41.1<sup>*</sup> | 41.2<sup>*</sup> |
| <sup>∞</sup>IP | — | — | 37.2<sup>**</sup> | 47.6<sup>*</sup> | 15.4<sup>**</sup> |
| Macroinvertebrates |  |  |  |  |  |
| Lotic | 90.5<sup>*</sup> | 9.5<sup>*</sup> | 27.7<sup>*</sup> | 46.5<sup>*</sup> | 25.8<sup>*</sup> |
| §BH wando | 97.1<sup>*</sup> | 2.9<sup>*</sup> | 15.4<sup>**</sup> | 67.9<sup>**</sup> | 16.7<sup>**</sup> |
| ¶BT wando | 88.2<sup>*</sup> | 11.8<sup>*</sup> | 10.4<sup>**</sup> | 58.5<sup>**</sup> | 31.1<sup>*</sup> |
| ∞IP | — | — | 9.1<sup>**</sup> | 73.6<sup>**</sup> | 17.1<sup>**</sup> |

<sup>3</sup>
<sup>3</sup> †SFPOM, suspended fine particulate organic matter; ‡BFPOM, benthic fine particulate organic matter; §BH, bar-head; ¶BT, bar-tail; ∞IP, isolated pond. \*, \*\* $P < 0.05$ , Tukey–Kramer test.

For stable isotope analysis of macroinvertebrates, we selected 91 taxa with > 0.2 mg dry mass. The δ^13^C and δ^15^N values of macroinvertebrates fell within the habitat-specific ranges among the three potential food sources (epilithon, leaves and dam plankton) in the dam outlet reaches. The trophic contributions of dam plankton to the macroinvertebrate community estimated by the three-source mixing model were significantly different among the four habitat types (*F* _3, 397_ = 23.462, *P* < 0.001) and among FFGs (*F* _4, 396_ = 35.695, *P* < 0.001). Lotic habitats had a significantly higher contribution from dam plankton (mean, 27.7%) than did the BH (15.4%), BT (10.4%) and IP (9.1%) (*P* < 0.001, Tukey–Kramer test). The contribution of leaves in the lotic and BT habitats was significantly higher than that in the BH and IP in the dam outlet reaches (*P* < 0.05, Tukey–Kramer test). Epilithon was estimated to be a principal food source throughout the habitat types downstream from dams.

In dam outlet reaches, filter feeders consumed significantly more dam plankton than the other FFGs throughout all habitat types (*P* < 0.001, Tukey–Kramer test). The contribution of dam plankton to filter feeders was higher in the lotic habitat than in the lentic habitats (*P* < 0.001, Tukey–Kramer test with Bonferroni correction), whereas the significant differences were absent for the other FFGs (Fig. 1). In lotic habitats in dam outlet reaches, spatial variation in the contribution of dam plankton to filter feeders was significantly positively correlated with the proportion of dam plankton in SFPOM (Fig. 2), whereas this correlation was not found for lotic habitats and the other FFGs.

**Figure 1.**
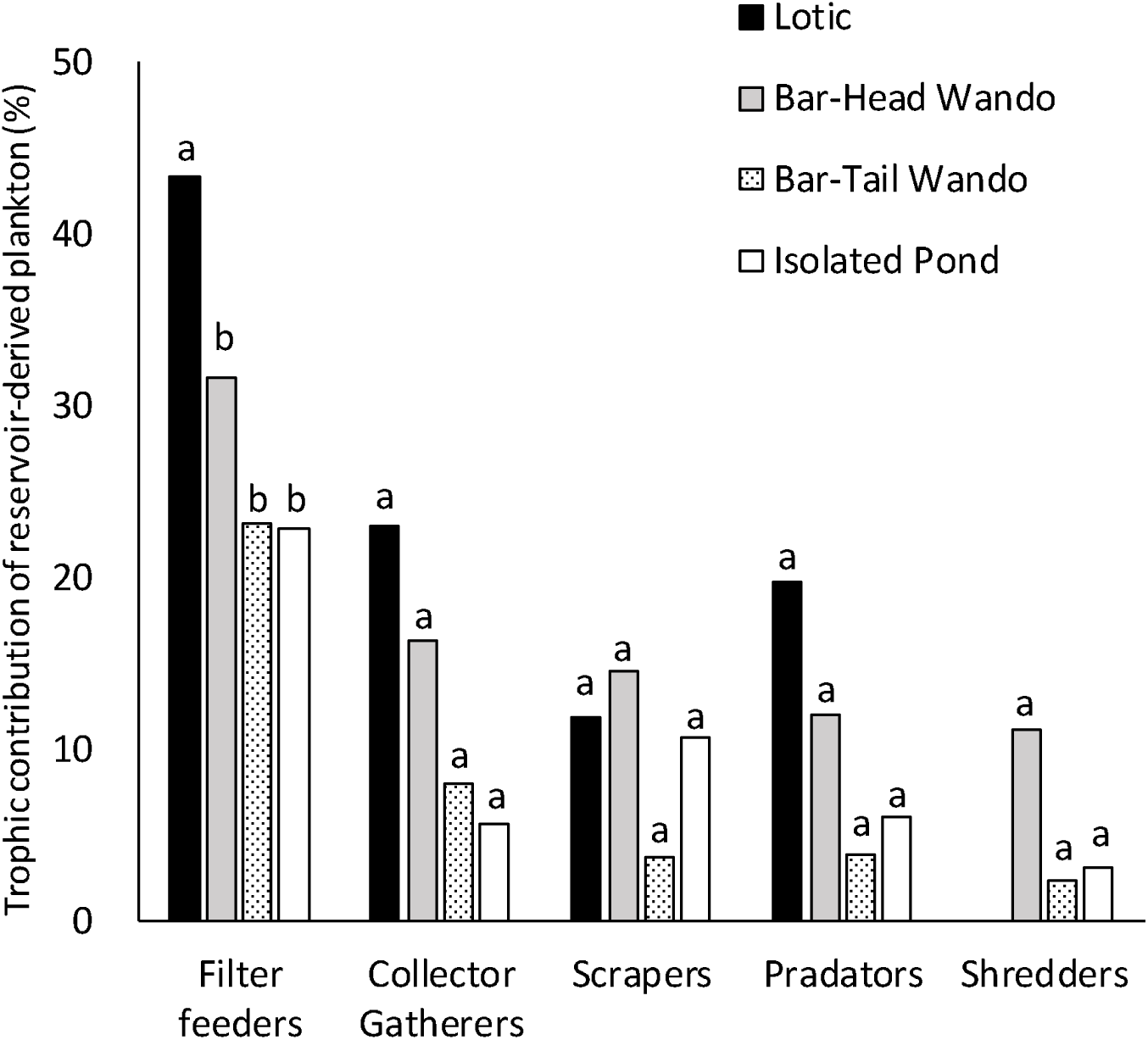
Comparison of trophic contributions of reservoir-derived plankton to each functional feeding group (FFG) among habitat types. Letters indicate significant differences among habitat types (P < 0.001, Tukey–Kramer test with Bonferroni correction).

**Figure 2.**
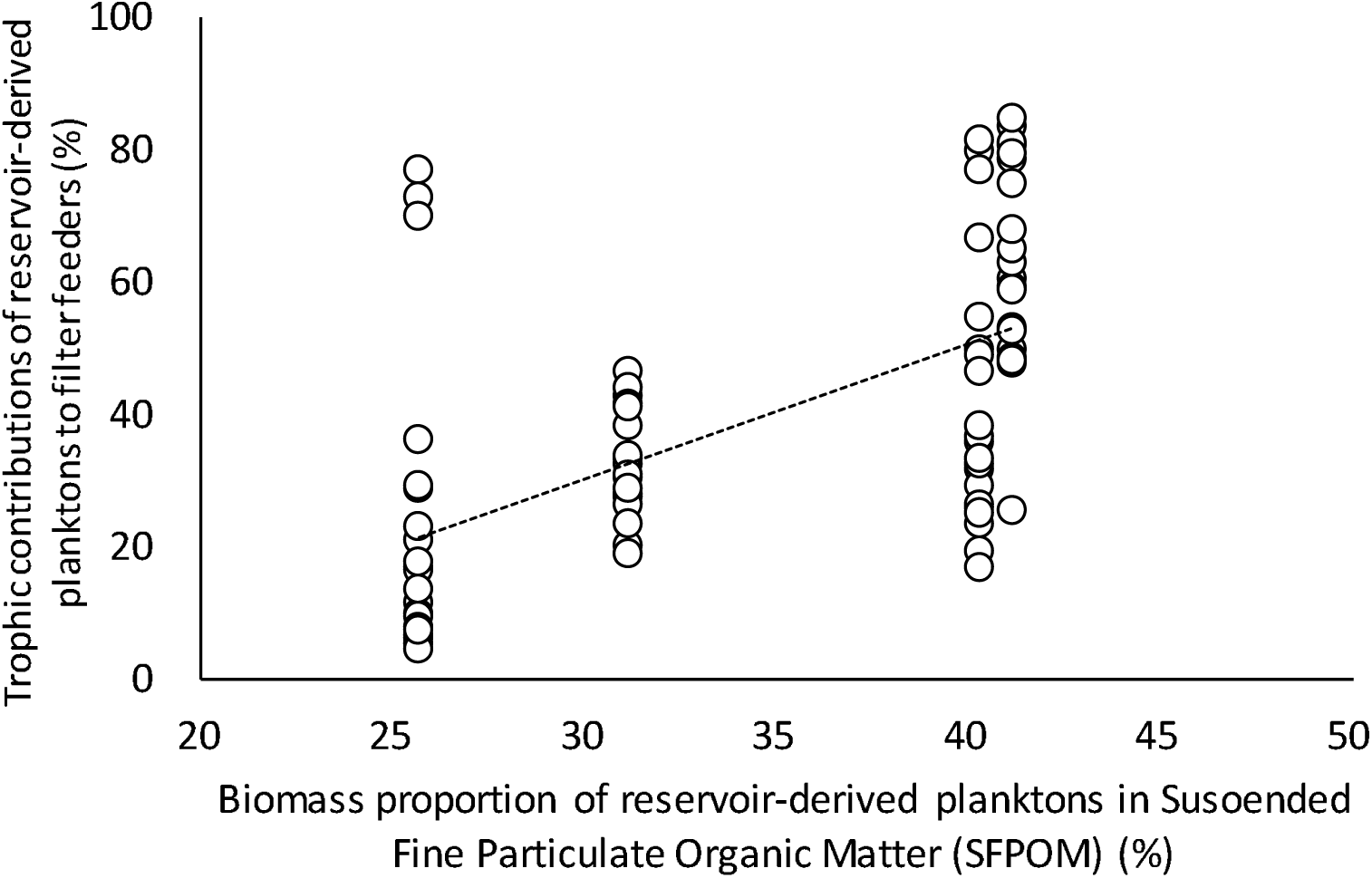
Relationships between trophic contributions of reservoir-derived plankton to filter feeders and percentage of reservoir-derived plankton in suspended fine particulate organic matter (SFPOM) in the lotic habitat. The regression line indicates significant correlation found in the lotic habitat (*r* = 0.574; *P* < 0.001). The other functional feeding groups (FFGs) while this correlation was not true for lotic habitats and the other FFGs.

## Discussion

Our approach to comparing trophic structures among different habitats has provided useful insights into the role of spatial environmental heterogeneity in the formation of food web structures in rivers. Environmental heterogeneity was related to community diversity and food resources of macroinvertebrates on each spatial scale (Boyero, 2003; Boyero & Bosch, 2004; Zilli & Marchese, 2011; Pilotto et al., 2016). However, in previous studies, the relationship between heterogeneity and food webs in macroinvertebrates within habitats was not considered. Using carbon and nitrogen stable isotope analyses, we compared the trophic influence of reservoir-derived plankton on downstream macroinvertebrate communities between lentic and lotic habitats. Our overall results showed habitat-specific patterns of trophic influence of dam plankton, with less influence of dam plankton in lentic than in lotic habitats. This pattern was specifically notable for filter feeders that feed on SFPOM. Although the high trophic influence of dam plankton and dam-derived FPOM on filter feeders has been reported in numerous studies (Doi et al., 2008; Power et al., 2013; Tagliaferro et al., 2013; Four et al., 2019), these earlier observations were limited to lotic habitats, where the strongest impact was observed in this study.

We considered the habitat-specific source composition of SFPOM as a potential driver of the reduced trophic impact in lentic habitats. The three-source model analysis of SFPOM indicated a lower proportion of dam plankton and higher proportions of epilithon and leaves in lentic habitats than in lotic habitats. The lower proportion of dam plankton in lentic SFPOM was most likely reflected in the lower trophic impact on lentic consumers through assimilation of SFPOM, especially on filter feeders. This trophic reflection of drifting SFPOM to filter feeders at dam outlet reaches is supported by previous isotopic studies (Doi et al., 2008; Wellard Kelly et al., 2013), although the data from lotic habitats are limited. The habitat-specific composition of SFPOM may be derived from hydraulic and landscape characteristics of lentic habitats. BH located at the leading edges of the bar was not only an area where surface water downwelled into the hyporheic zone but also an area where SFPOM from upstream was accumulated (Boulton et al., 2008). Hence, the contribution of dam plankton in the BH was higher than that in other lentic habitats, but the amounts of epilithon and leaves were higher than in the main stream, resulting in an increase in river-derived SFPOM. It can be inferred that the impact of the dam was moderated. BT located at the downstream end of the bar area where groundwater upwells is an environment where mainstream water and SFPOM cannot easily enter. The biomass of organic matter in the BT is also higher than in the mainstream, and the interaction between hydromorphological and ecological processes may have contributed to mitigating dam effects. On the other hand, it is impossible for dam plankton to flow into the IP when it is disconnected from the mainstream during periods of normal water level. After a flood, dam plankton may be left behind in the IP. In addition, the amount of BFPOM in the IP may have increased as a result of plankton production in the water body (Doi, 2009). Therefore, it was suggested that the influence of dam-derived plankton in lentic habitats is due to different hydromorphological and ecological processes. Furthermore, SFPOM derived from allochthonous and autochthonous sources was produced by decomposition of bacteria and relatively dominant scrapers and shredders that inhabited the lentic habitat (Langhans et al., 2008; Treplin & Zimmer, 2012; Halvorson et al., 2015). The high amounts of epilithon, BFPOM and BCPOM observed in lentic habitats can account for these geohydraulic and biological processes, leading to the higher proportions of epilithon and leaves in lentic SFPOM.

In addition to the habitat-specific composition of food sources, habitat preference of macroinvertebrates was considered as a potential reason for the reduced trophic contribution of dam plankton in lentic habitats. Filter feeders generally prefer to live in lotic environments with rapid flow and loose stones and gravel of suitable size for their net-spinning behaviour on the riverbed (Georgian & Thorp, 1992). Stabilisation of the substrate due to reduced hydraulic and sediment dynamics resulting from dam control is also a driving factor for the abundance of filter feeders at dam outlet reaches (Oswood, 1979; Hoffsten, 1999; Tszydel et al., 2009). In lentic habitats, the substrate is mainly composed of fine materials (silt and sand) and embedded stones with extremely low water flow. Therefore, the abundance ratio of filter feeders in the macroinvertebrate community is reduced in lentic habitats, with increases in the ratios of other FFGs, such as collector-gatherers, scrapers and shredders (Table 2). These other FFGs can feed on sources less mixed with reservoir-derived plankton (e.g. BCPOM and BFPOM) and thus are robust to the high input of dam plankton. As a result of high ratios of these FFGs being potentially robust to the input of dam plankton, the trophic impact on the macroinvertebrate community as a whole may be reduced.

Finally, from the conservation viewpoint, the ecologically important role of lentic habitats is worth mentioning. Although the mean proportion of lentic area in the downstream reaches was less than 5.7% in our study, the distinctive hydro-physical and landscape characteristics of the lentic areas led to heterogeneous trophic conditions within the reaches. In general, the amount of lentic habitat in sand and gravel bars decreased in downstream reaches due to riverbed degradation (Brandt, 2000; Rollet et al., 2013). In this study, the area of lentic habitat was lower in downstream reaches than in upper reaches, except for KD3. Because KD3 was the most distant reach from the dam, it was believed that the lentic habitat was restored by sediment supplied from the river bank and reduction of the impact of the dam (Rollet et al., 2013). In previous studies, the significance of lentic habitats in the formation of biodiversity was often reported (Taniguchi & Tokeshi, 2004; Tews et al., 2004; Warfe et al., 2008) and was applied to conservation. For example, many river restoration works have created lentic spaces to facilitate spatial heterogeneity of species diversity (Wyżga et al., 2012; Van den Brink et al., 2013) and to supply flow refugia to fishes from flood disturbances (Sedell et al., 1990; Milner & Gilvear, 2012). Creation of lentic habitat was basal idea downstream due to improvement of river morphology, and it was shown that it contributed to mitigation of the trophic impact. Our finding of habitat-specific food resources in lentic spaces and their influence on trophic structures of invertebrate communities expanded the existing idea a little further. Lentic water bodies can help sustain adequate and diverse flows of materials in food webs. In particular, in environments where the main food sources are artificially modified from the natural sources, such as in the dam outlet reaches observed in this study, lentic habitats are expected to act as trophic refugia that can mitigate the trophic impact.

## Acknowledgments

This research was supported by the Japan Society for the Promotion of Science (grant numbers 25241024 and 26257304). We thank Sakiko Yaegashi, Shoichi Suzuki and Yukihiro Kumagai for the assistance with fieldwork and laboratory analysis.

## Data Availability Statement

The data that support the findings of this study are available from the corresponding author upon reasonable request.

## Conflict of Interest Statement

The authors declare that they have no competing interests.

---

**Figure S1.**
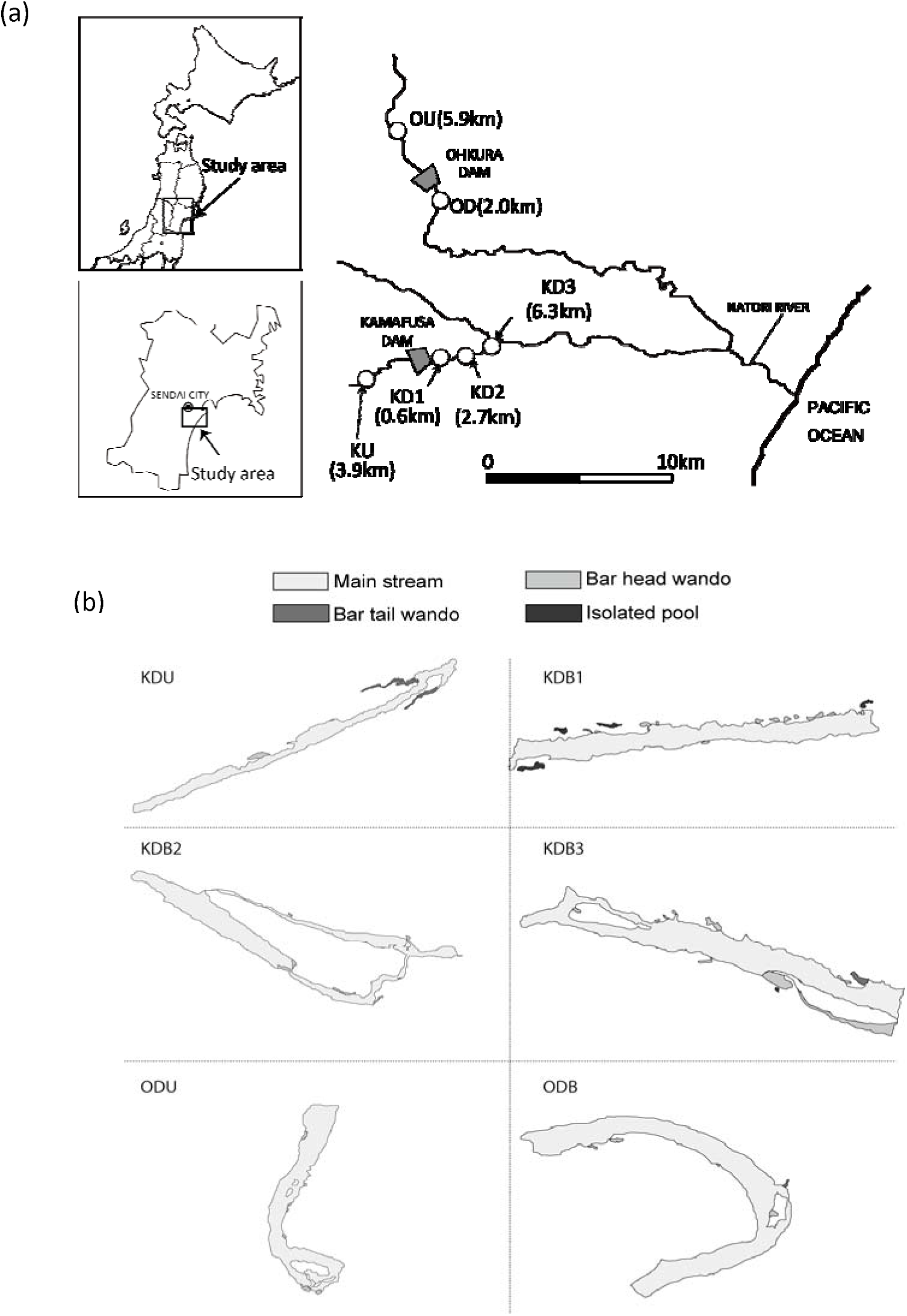
Maps of study site (a) and habitats (b). KU, Kamafusa dam upstream; KD1–KD3, Kamafusa dam downstream 1–3; OU, Ohkura dam upstream; OD, Ohkura dam downstream.

